# Revisiting the paradigm of anhematophagy in male mosquitoes

**DOI:** 10.1101/2024.10.08.617226

**Authors:** Jovana Bozic, Renuka E. Joseph, Tse-Yu Chen, Margaret V. Becker, Rachel S. Krizek, Amber Holley, Renee L.M.N. Ali, Miguel Á. Miranda, Carlos Barceló, Mikel A. González, Maureen Laroche, Rafael Gutiérrez-López, Renke Lühken, Douglas E. Norris, Perran A. Ross, Joshua B. Benoit, Jason L. Rasgon

## Abstract

Most female mosquitoes are reproductively obligate feeders on vertebrate blood to obtain nutrients required for egg production (driving transmission of vector-borne pathogens in the process), and which rely on plant sugars for their non-reproductive energy requirements. Male mosquitoes, on the other hand, are thought to rely exclusively on plant sugars for their energetic needs; indeed, this dichotomy is one of the central paradigms of medical entomology. Here, we provide multiple lines of evidence challenging this assumption. We show that when reared under dehydration/starvation conditions, male mosquitoes will readily take blood from a membrane feeder while CRISPR mutants with impaired humidity detection do not increase their bloodfeeding rates. For multiple mosquito species, dehydrated males are attracted to human hosts and attempt to probe. We also observed several instances where dehydrated probing males were able to successfully pierce vertebrate host skin and acquire blood from a vertebrate host. When fed a blood meal containing West Nile virus, male *Cx. tarsalis* mosquitoes became infected with and orally transmitted the pathogen at rates and titers equivalent to females. Finally, we collected wild, putatively blood-fed male mosquitoes from Texas, USA and Mallorca, Spain, and identified vertebrate DNA in these samples (canine and human, respectively). These data suggest that under specific circumstances male mosquitoes may be able to probe and/or ingest blood and transmit pathogens to vertebrate hosts. These results are a paradigm shift in our understanding of male mosquito biology and suggest they may be more directly involved in pathogen transmission cycles than previously recognized.

## Introduction

Female mosquitoes (except those which are autogenous) are reproductively obligate bloodfeeders, feeding on vertebrate blood to obtain nutrients required for egg production, and relying on plant sugars for their non-reproductive energy requirements. In contrast, male mosquitoes, on the other hand, are thought to rely exclusively on plant sugars for their energetic needs [1]. Indeed, this difference is one of the central tenants of medical entomology; female mosquitoes bloodfeed, males do not. Female bloodfeeding facilitates transmission of blood-borne pathogens, such as viruses or parasites, between vertebrate hosts, which is why the majority of mosquito research is performed on females rather than males; even when males are studied, it is usually within the context of how they affect females (mating behavior and fertility, pathogen transmission modulation, etc.) [2–5].

Female mosquitoes evolved to feed on blood, and this adaptation is reflected in the biology of their midgut, where transcripts related to blood digestion are enriched in the female compared to the male [6]; one would expect that male mosquitoes should not be attracted to blood as a nutrition source as they are thought to lack the proper physiology to digest and process it. However, there is one interesting report in the literature where male mosquitoes were attracted to and fed on blood. Nikbakhtzadeh and colleagues [7] documented bloodfeeding behavior in a laboratory colony of the mosquito *Culex quinquefasciatus.* When presented with defibrinated sheep blood on a cotton pledget (and to a much less efficient extent, a Parafilm membrane), male mosquitoes took a bloodmeal. However, in this study blood was toxic to male mosquitoes, which died in a dose-dependent manner when blood was mixed with sugar [7], consistent with physiological adaptations to sugar vs bloodfeeding in males vs. females [6]. Interestingly, males did not show a preference for sugar compared to blood in a dual-choice assay [7], and the reason they fed on blood at all, particularly as it was toxic, remains an open question. As this is (to our knowledge) the only published observation of male mosquito bloodfeeding behavior, it is difficult to speculate. However, there are a multitude of observations that male mosquitoes are attracted to human host odors and this behavior is suppressed by mosquito repellents [8], which includes species from arguably the three most important mosquito genera that act as pathogen vectors to humans (*Anopheles*, *Culex*, and *Aedes*).

Here, we present studies on male bloodfeeding behavior in the mosquitoes *Cx. tarsalis, Cx. quinquefasciatus, Ae. aegypti, Ae. notoscriptus,* and *An. stephensi*. *Cx. tarsalis* is one of the major West Nile virus (WNV) vectors in North America, where it is widely distributed across the western United States [9]. It is genetically diverse, generally feeds on birds in the wild, and can be facultatively autogenous [9–10]. After becoming infected with WNV during a bloodfeeding event, it can also transmit the virus vertically to offspring at relatively high rates [11–12]. *Ae. aegypti* is one of the major invasive arbovirus vectors in the world [13]. *An. stephensi* is a major vector of malaria in Asia, and has been recently expanded its range into Africa [14]. *Cx. quinquefasciatus* is a major arbovirus vector with a cosmopolitan distribution [15], while *Ae. notoscriptus* is a vector of Ross River virus in Australia [16]. We opportunistically observed *Cx. tarsalis* and *Ae. aegypti* males taking blood during unrelated laboratory studies, and undertook experiments to document and understand the behavior in multiple species. We found that when dehydrated, male mosquitoes will predictably take blood from a membrane feeder, determined the physiological mechanisms driving male bloodfeeding behavior, demonstrated that some males were capable of obtaining a bloodmeal from a vertebrate host through biting, and present results of experiments examining potential for male mosquitoes to be involved in pathogen transmission cycles. We also collected bloodfed male mosquitoes in the wild and identified the hosts that they fed upon. These results are a paradigm shift in our understanding of male mosquito biology and suggest they may be more directly involved in pathogen transmission cycles than previously recognized.

## Methods

### Human and animal subjects approvals

Human landing experiments with *Cx. tarsalis* used the senior author (JLR) under PSU IRB Exempt Protocol STUDY00024284. Human landing experiments with *Aedes* mosquitoes and *Cx. quinquefasciatus* used author PAR under University of Melbourne Human Ethics committee approval 0723847. Experiments with mice were conducted under JHU IACUC protocol MO21H10.

### Mosquitoes

*Cx. tarsalis* (KNWR strain), *Cx. quinquefasciatus* (sourced from Queensland, Australia), *Ae*. *aegypti* (Liverpool, IR93a mutant, or Cairns strains), *Ae. notoscriptus* (sourced from Queensland, Australia), and *An. stephensi* (Liston strain) were maintained at 25°C, 16:8 h light:dark diurnal cycle with 80% relative humidity, with 10% sucrose solution provided at all times through a cotton wick. For general rearing, mosquitoes were provided with expired anonymous human blood through a water-jacketed glass membrane (Parafilm) feeder or a Hemotek feeder for egg development.

### Male dehydration

To stimulate bloodfeeding, male mosquitoes were held at 25°C, 75% RH without sugar or water for 24 hours [17].

### Survival analysis

Bloodfed male *Cx. tarsalis* mosquitoes were isolated and placed into cup cages, held at the previously described standard insectary conditions, and provided with a cotton ball soaked in 10% sucrose solution. Control males were non-bloodfed and were maintained under the same conditions. Dead mosquitoes were counted every day and removed from the cages. Significant differences in survival between mosquito groups was determined with Kaplan-Meier analysis using GraphPad Prism version 9.0.4.

### Dehydration and bloodfeeding behavior in male *Cx. tarsalis*

*Cx. tarsalis* males were reared conventionally (80% RH, with free access to 10% sucrose solution in water), or under dehydrating conditions as described above, then were offered a bloodmeal through a membrane feeder for 30 minutes. The number of fed and unfed mosquitoes at the end of the feeding period were counted. Data were analyzed by Fishers Exact test.

### Ionotropic receptor 93a (Ir93a) mutant mosquito assays

We obtained an *Ae. aegypti* line that was a CRISPR knockout mutant for the Ir93a gene, which inhibits their ability to sense humidity [18]. The mutation was introgressed into the wild-type Liverpool background for comparison with Liverpool controls, and both lines reared as described above. For experiments, at 5-6 days post-emergence, males of each strain were transferred to 10 x 10 x 10 cm cages and deprived of sucrose and water (or held at normal conditions as controls) for 24 hours before being offered an anonymous human bloodmeal (Innovative Research) using an artificial feeding system (Hemotek). Bloodfeeding rates for each genotype and condition were recorded. Data were analyzed by Fishers Exact tests and confidence intervals calculated from the binomial distribution.

### Landing and probing experiments

Cages of 50 male mosquitoes (*Cx. tarsalis*, *Cx. quinquefasciatus*, *Ae. notoscriptus,* or wild-type *Ae. aegypti*) were reared under standard or dehydrating insectary conditions and were allowed to probe on the hand of JLR (*Cx. tarsalis*) or PAR (*Ae. aegypti, Ae. notoscriptus,* and *Cx. quinquefasciatus*) for five minute trials. Mosquito landings (defined as a mosquito alighting on the volunteer hand for any period of time) and probing behavior (defined as exploring and probing with mouthparts) were counted during the 5 minute interval. Each experiment was repeated 6 times. Data were analyzed by Mann-Whitney U test.

### Host bloodfeeding by male mosquitoes

#### Cx. tarsalis

The senior author (JLR) had an unrelated small (3mm) scratch on their hand obtained from a pet cat a day earlier. A sterile razor blade was used to pick the scab off the scratch allowing a minimal amount of blood to be exposed. The wounded hand was placed in a cage of 20 dehydrated male *Cx. tarsalis* mosquitoes and their behavior recorded.

#### An. stephensi

Mosquitoes were reared at 27°C (± 1°C) with 75% ± 5% relative humidity and a 12-h day–night cycle by Johns Hopkins Malaria Research Institute Insectary Core facility. Male mosquitoes were transferred to cups (40-50 per cup) and held under dehydrating conditions for 24h. Prior to the feed the number of surviving mosquitoes was counted. Mosquitoes were allowed to feed on a sedated 8 week old female Swiss Webster mouse for 15 mins. The mouse was placed belly down on the top of the cup. No wound was created, and the fur was not shaved. Cups were immediately frozen after the attempted feed and mosquitoes screened under a microscope for the presence of a blood meal.

### WNV feeds

Dehydrated *Cx. tarsalis* males and female controls were allowed access to an infectious blood meal consisting of a 1:1 mix of anonymous human blood spiked with 5.0 x 10^7^ FFU/ml (focus-forming units/ml) of WN02-1956 (GenBank: AY590222). A subset of male and females were processed immediately after feeding (“day zero”) to check for virus viability. Mosquito virus infection and transmission assays were performed at 7 and 14 days post-blood feeding. Fully engorged mosquitoes were sorted from non-fed ones for analysis. Mosquitoes were anesthetized with triethylamine (Sigma, St. Louis, MO), legs/wings from each mosquito were removed and placed separately in a 2-mL tube filled with 0.5 mL mosquito diluent (MD: 20% heat-inactivated fetal bovine serum (FBS) in Dulbecco’s phosphate-buffered saline, 50 µg/mL penicillin/streptomycin, 50 ug/mL gentamicin, and 2.5 µg/mL fungizone, with a sterile 2.0 mm stainless steel bead (Next Advance, Inc. Innovative Lab Products for the Life Sciences). The proboscis of each mosquito was positioned in a tapered capillary tube containing approximately 10 µL of a 1:1 solution of 50% sucrose and FBS to induce salivation. After 30 min, the tube contents were expelled into 0.3 mL MD, and bodies were placed individually into a 2-mL tube filled with 0.5 mL MD and a stainless steel bead as described above. Mosquito bodies and legs/wings were homogenized for 30 sec with TissueLyser (QIAGEN, Hilden, Germany) at 24 cycles/sec, followed by centrifugation for 1 min. Mosquito bodies, legs/wings, and salivary secretion samples were tested for live, infectious WNV using focus-forming assays (FFAs; see below).

### WNV FFAs

WNV titers were quantified by FFA, which detects live, infectious virus. C6/36 cells were seeded into 96-well plates at a density of 1X10^5^ cells/well and incubated overnight at 28°C in complete RPMI medium without CO_2_. The next day, medium was removed from the wells. Samples from male and female bodies or legs/wings were serially diluted in a serum-free RPMI medium; saliva samples were undiluted. 30 μL of each sample was added in duplicate to the prepared C6/36 cells.

Cells were incubated for 1 hour at 28°C without CO_2_, after which the inoculum was removed. 100 μL of RPMI containing 0.8% methylcellulose was added to limit viral spread. Infected cells were incubated for 48 hours at 28°C without CO_2_. At 48 hours post-infection, infected C6/36 cells were fixed with 50 μL 4% formaldehyde for 30 minutes at room temperature (RT). Cells were washed, permeabilized with 0.2% triton-X, and blocked with 3% BSA. 30 μL monoclonal flavivirus antibody (Clone D1-4G2-4-15, BEI-resources, NR-50327) was added and incubated overnight at 4°C. After washing, 30μl of fluorescent secondary antibody (1:1000 dilution; Goat anti-Mouse IgG (H+L) Highly Cross-Adsorbed Secondary Antibody, Alexa Fluor 488, Invitrogen/Thermo Fischer, A-11029) was added and incubated overnight at 4°C. Cells were maintained in 100 μL PBS to prevent drying. WNV foci were imaged using a FITC filter on an Olympus BX41 microscope with a UPlanFI 4x objective and counted. Infection rate (IR) was defined as the proportion of mosquitoes exposed to virus that had WNV-positive bodies. Dissemination rate (DR) was defined as the proportion of mosquitoes with WNV-positive bodies that had WNV-positive legs/wings. Transmission rate (TR) was defined as the proportion of mosquitoes with WNV-positive legs/wings that had WNV-positive saliva. IR, DR, and TR were analyzed with Fisher’s exact tests. Viral titers were analyzed using Mann-Whitney U tests.

### Wild mosquito collection, DNA extraction and molecular detection and identification of vertebrate DNA

Mosquitoes were collected using BG-Sentinel traps (Biogents) [19–20] on Galveston Island, in South Texas, August 12-13, 2024, and in Mallorca, Spain, July 9, 2020. Fresh mosquitoes were sexed, morphologically identified the same day and placed at -20°C until processed. For Texas samples, after identification, whole male mosquitoes were individually incubated with 20µL of proteinase K and 180µL of G2 buffer (Qiagen) at 56°C for one hour then were homogenized using tungsten beads and a TissueLyser (Qiagen) for one minute at 30 cycles/sec. After a quick spin to pellet debris, DNA was extracted from the lysate using a Qiagen EZ1 automated DNA extraction instrument and reagents from the associated EZ1&2 DNA Tissue Kit. DNA samples were screened for vertebrate DNA using vertebrate-specific qPCR assay targeting a fragment of the 16s rRNA on a CFX Duet PCR system (Bio-Rad) [21].

Human blood obtained from the University of Texas Medical Branch Blood Bank was used as a positive control and PCR mix without DNA was used as a negative control. Samples were analyzed in duplicate and the entire assay repeated twice. Samples positive for vertebrate DNA had a 690 bp sequence of the COI gene amplified by primers ModRepCOIF and ModRepCOIR [22], the amplicons cloned using the NEB PCR cloning kit (New England Biolabs, Ipswich, MA), sequenced, and identity confirmed by BLAST analysis. For Mallorca samples, whole engorged specimens were placed into a 2 mL tube, along with 20 2.0 mm zirconia beads (BioSpec Products, Bartlesville, USA) and 1 mL of cell culture medium (high-glucose Dulbecco’s modified Eagle’s medium; Sigma-Aldrich, St. Louis, MO, USA) and homogenized with a TissueLyser (Qiagen, Hilden, Germany) for 2 minutes at 50 oscillations per second. Following clarification through centrifugation for 1 minute at 8000 rpm and 4°C, the resulting suspension was transferred to a new safe-lock tube. DNA was extracted from 200 μL of the homogenate using the KingFisher™ Flex Magnetic Particle Processor with the MagMAX™ Pathogen Ribonucleic Acid/DNA Kit (both Thermo Fisher Scientific, Waltham, MA, USA). A primer set targeting the 16S rRNA gene was used following previously published protocols [23]. All amplicons were directly sequenced (LGC Genomics, Berlin, Germany).

Conceptual replication: As we are aware that the results presented here are controversial, we endeavored to have multiple independent laboratories conduct the experiments to test the generality of the observations. Work with *Cx. tarsalis* was performed at Penn State; work with mutant *Ae. aegypti* was conducted at the University of Cincinnati; work with *Ae. notoscriptus, Cx. quinquefasciatus,* and wild-type *Ae. aegypti* was conducted at the University of Melbourne; work with *An. stephensi* was conducted at the Johns Hopkins Bloomberg School of Public Health.

## Results

### Bloodfeeding is not toxic to *Cx. tarsalis* males

While we were bloodfeeding during an experiment related to relative humidity (e.g. [24]), we noted incidentally that in addition to females, male mosquitoes were probing the membrane and were taking blood (Figure 1A,B). As this was a spontaneous occurrence, the total number of males in the cage was not recorded but was on the order of 50-70 based on standard rearing practices in our lab. Out of this total, we isolated seven blood-engorged males. These males were placed into a survival cup and a survival experiment conducted, comparing their survival to 10 non-bloodfed males from the same initial cage. Although Nikbakhtzadeh et al. [7] demonstrated that blood was highly toxic to male *Cx. quinquefasciatus* in laboratory studies, we did not observe any acute toxicity to blood in male *Cx. tarsalis*; indeed, survival in bloodfed males was marginally (although not statistically) higher than non-bloodfed males (Figure 1C).

**Figure 1.**
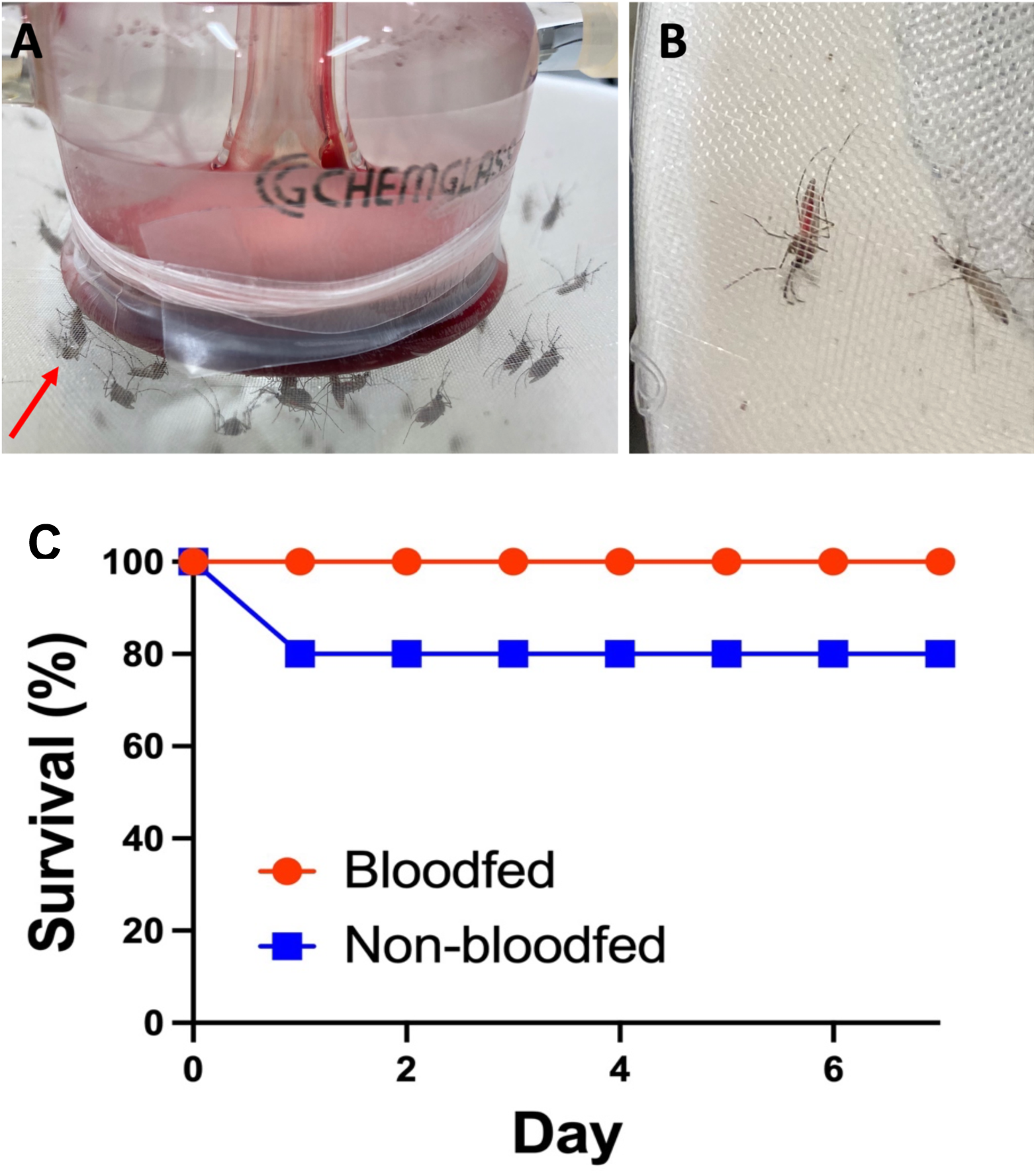
Male *Cx. tarsalis* bloodfeeding behavior and survival. A) mosquitoes congregating at and bloodfeeding from a paraffin membrane. Arrow points to male orienting toward the membrane. B) Blood-engorged male mosquito. Unengorged male can be seen in-frame. C) Survival curve of bloodfed vs. non-bloodfed male *Cx. tarsalis* mosquitoes. No significant difference was observed between treatments.

### *Cx. tarsalis* male bloodfeeding is driven by dehydration

As we previously demonstrated that dehydration stimulates elevated bloodfeeding behavior in females [17, 24–25], we tested the hypothesis that dehydration was driving bloodfeeding behavior in males. Cages of male mosquitoes were reared under conventional insectary conditions or under dehydrating conditions [17]. No conventionally reared male mosquito (N = 64) took a bloodmeal from the membrane feeder, while 44/163 dehydrated males took a bloodmeal (*P* < 0.00001).

### Male mosquito bloodfeeding behavior is dependent on their ability to sense humidity

Mosquitoes sense humidity through ionotropic receptor Ir93a, by which they locate oviposition sites, and CRISPR Ir93a knock-out mutants are impaired in this behavior [18]. We noted in further experiments that male *Ae. aegypti* mosquitoes would also take blood from a membrane feeder, so we used an available Ir93a *Ae. aegypti* KO mutant for these assays [18]. When reared under standard insectary conditions, bloodfeeding rates did not differ statistically between wild-type and mutant mosquitoes. However, when reared under dehydrating conditions, bloodfeeding rates for the mutant did not increase, while wild-type mosquitoes had significantly elevated bloodfeeding behavior (*P* = 0.0284) (Figure 2).

**Figure 2.**
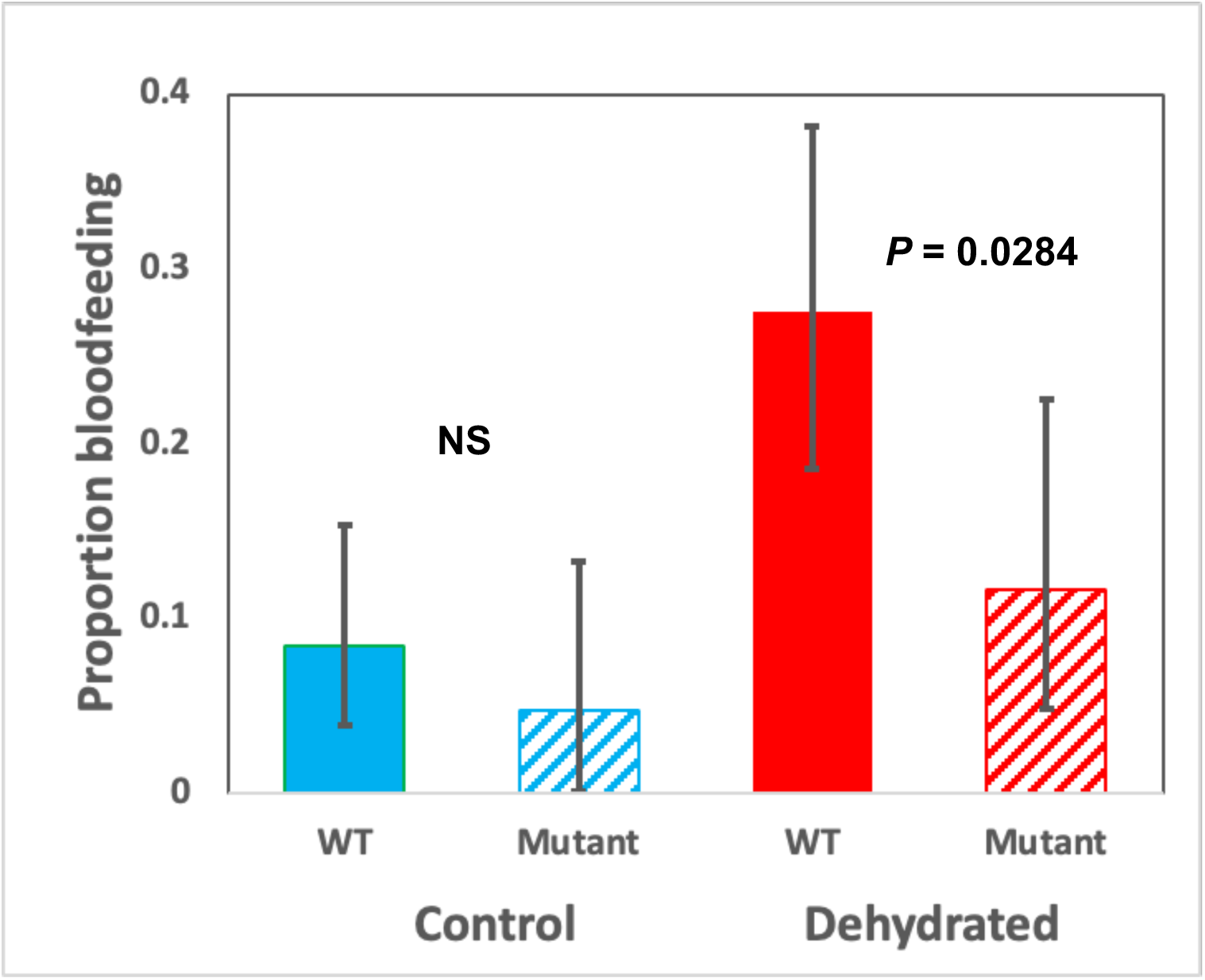
CRISPR deletion of Ir93a ablates male mosquito bloodfeeding behavior under dehydration conditions. When reared conventionally, both wild-type and mutant *Ae. aegypti* exhibit baseline levels of bloodfeeding behavior. When reared under dehydration conditions, wild-type males significantly increase bloodfeeding behavior but humidity-insensitive mutant mosquitoes do not. Confidence intervals were calculated from the binomial distribution. WT = wild-type.

### Dehydrated male mosquitoes will probe the hand of a human volunteer

The hand of a human volunteer was exposed to cages of conventionally reared or dehydrated male mosquitoes (*Cx. tarsalis, Cx. quinquefasciatus, Ae. aegypti, and Ae. notoscriptus*). Conventionally reared males showed little interest in the host, with infrequent landings that lasted less than 5 seconds. In all trials, only one conventionally reared mosquito (*Ae. aegypti*) demonstrated probing behavior. In contrast, for all tested species dehydrated males landed significantly more often on the hand of the volunteer, most landings lasted until the end of the time period, and probing behavior was observed in the majority of landings (Figures 3-4, Supplementary Videos 1-9). One dehydrated *Cx. tarsalis* male mosquito (out of 6 separate trials) was able to pierce the skin of the volunteer at the base of the wrist, although it was unable to reach the capillaries and acquire a bloodmeal (Figure 4, Supplementary Video 10). The bite resulted in a mild immunogenic reaction that disappeared after approximately 10 minutes (Figure 4G,H). To confirm that only males were in the cage, after the study was concluded the entire cage was killed by freezing and every mosquito visually examined for the presence of a female or a gynandromorph; only males were identified.

**Figure 3.**
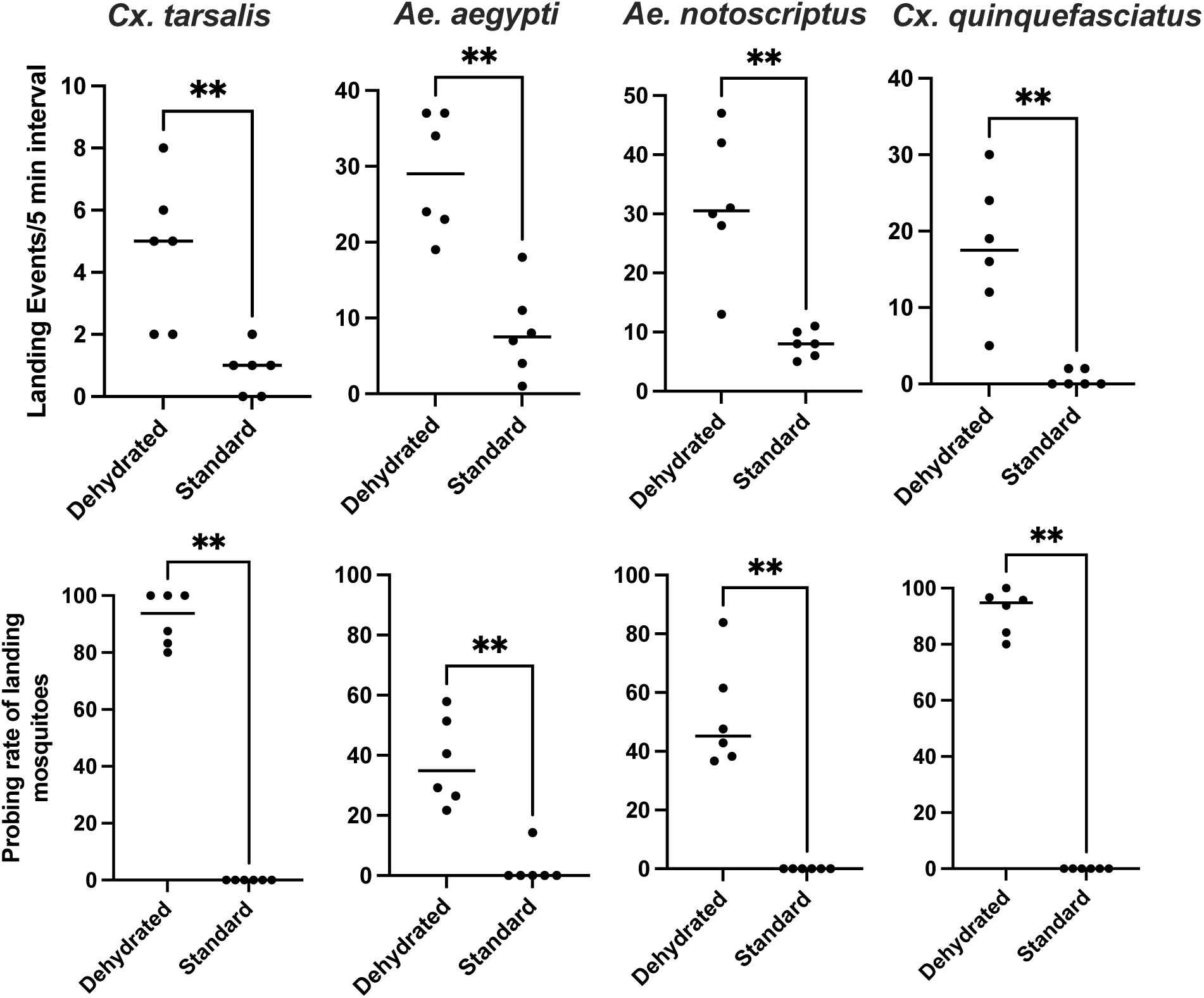
Host probing behavior of dehydrated male mosquitoes. Top: Landing responses for dehydrated vs. conventionally reared (“standard”) male mosquitoes. Bottom: Probing behavior for dehydrated vs. conventionally reared (“standard”) male mosquitoes. ** = *P* < 0.01

**Figure 4.**
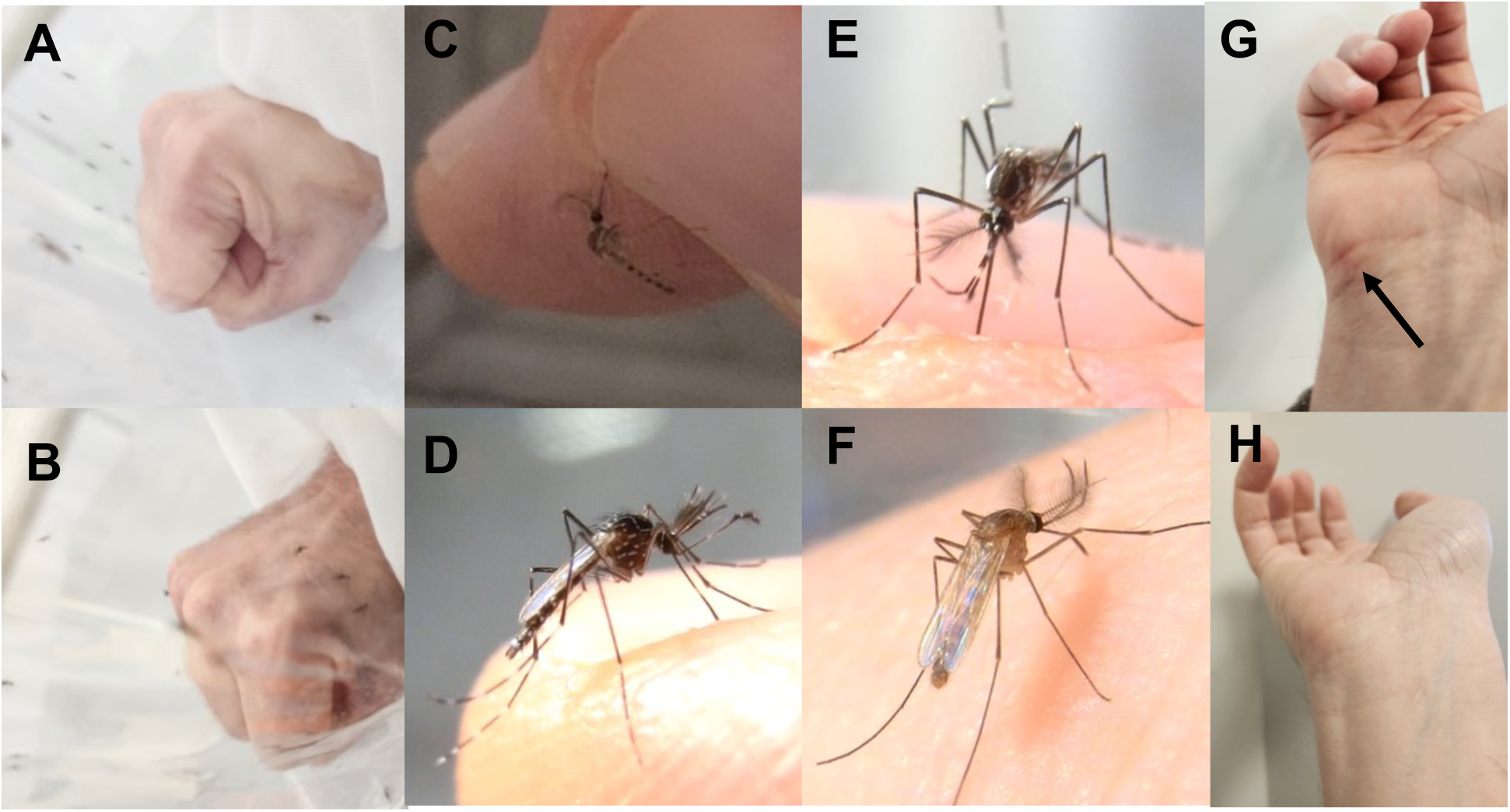
Landing and probing behavior of dehydrated mosquitoes. A) Conventionally reared mosquitoes (*Cx. tarsalis*) ignore a human host, while B) dehydrated mosquitoes actively land on a human host. Dehydrated mosquitoes C) *Cx. tarsalis* D) *Ae. aegypti* E) *Ae. notoscriptus* F) *Cx. quinquefasciatus* actively probe after landing. G) Bite reaction 2 minutes post-probing (arrow) H) Immune reaction resolved by 10 minutes post-probing. See Supplementary videos 1-9 for complete behavioral responses of each mosquito species and Supplementary video 10 for biting behavior.

### Dehydrated male mosquitoes will take blood from vertebrate hosts

Dehydrated male mosquitoes show keen interest in probing a human host, but in these experiments were not able to acquire blood, even from the single observed “successful” probing attempt. We hypothesized that if blood was made more accessible, male mosquitoes would take a bloodmeal. The senior author serendipitously had a small scratch on their hand (acquired from a pet cat a day earlier). The scab was peeled back using a sterile razor blade, exposing a small amount of blood. The volunteer placed their hand in a cage of 20 dehydrated *Cx. tarsalis* male mosquitoes. Males were attracted to the wound, and wound probing behavior was observed by 5 males (Supplementary Video 11). One male out of the 5 that probed fed and took a bloodmeal from the wound (Figure 5A-C). At the conclusion of the experiment, the fed male was dissected to confirm the presence of blood in the gut (Figure 5D).

**Figure 5.**
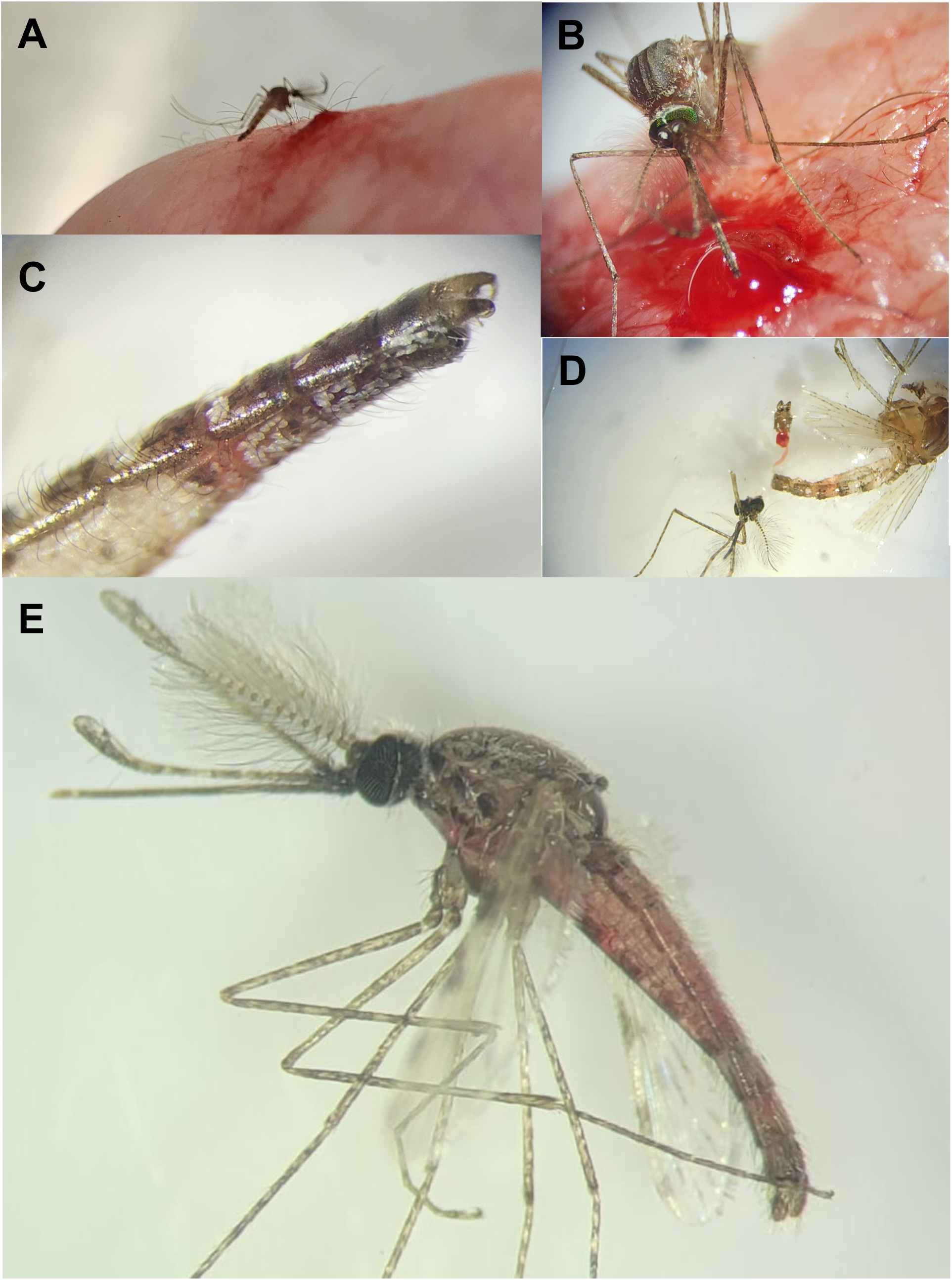
Male mosquito bloodfeeding. A) Male *Cx. tarsalis* feeding on an open wound. B) Close-up of *Cx. tarsalis* feeding behavior. C) Blood is observable in the male *Cx. tarsalis* mosquito gut. D) Blood in the male *Cx. tarsalis* mosquito gut was confirmed by dissection. E) Male *An. stephensi* mosquito with visible blood in the midgut after feeding on a mouse. See Supplementary Video 11 for complete *Cx. tarsalis* behavioral response.

Further experiments were conducted examining the ability for male *An. stephensi* mosquitoes to take blood from a mouse host (which has thinner skin compared to a human). *An. stephensi* males were more sensitive to the dehydration procedure compared to *Aedes* or *Culex*, and out of the 2 replicate experiments most did not survive the dehydration procedure (Rep 1: 34/39 dead; Rep 2: 46/50 dead). However, out of the 9 surviving mosquitoes across both replicates, one mosquito (in replicate 1) was able to bite the mouse and acquire a bloodmeal (Figure 5E). These data prove that male mosquitoes do have the ability to pierce the skin of some vertebrates and can canulate capillaries to acquire blood.

### Male *Cx. tarsalis* mosquitoes are competent vectors for West Nile virus

Since we determined that male mosquitoes will probe vertebrate hosts, can pierce skin, and ingest blood, we asked the question: can male mosquitoes become infected with and transmit pathogens? We offered dehydrated male *Cx. tarsalis* mosquitoes an artificial bloodmeal spiked with WNV through a membrane feeder and assayed their virus vector competence at day 7 and day 14 post-feeding. Female *Cx. tarsalis* were exposed to virus at the same time as a control. We found that both female and male *Cx. tarsalis* were able to become infected with, disseminate, and orally transmit virus; males transmitted at both day 7 and 14, while females only had detectable virus in their saliva at day 14. After adjusting for multiple comparisons, infection rates (IR), dissemination rates (DR), and transmission rates (TR) did not differ statistically between males and females at either timepoint (Table 1).

**Table 1.** WNV Infection rate (IR), dissemination rate (DR) and transmission rate (TR) of dehydrated males and females at 7 and 14 days post-infection. No comparisons were statistically significant after correcting for multiple comparisons.

|  | N | IR | DR | TR |
| --- | --- | --- | --- | --- |
| Day 7 |  |  |  |  |
| Males | 43 | 0.53 | 0.70 | 0.56 |
| Females | 10 | 0.50 | 0.60 | 0.00 |
| Day 14 |  |  |  |  |
| Males | 14 | 0.50 | 0.86 | 0.50 |
| Females | 12 | 0.92 | 0.91 | 0.30 |

We quantitated all viral titers using an infectious virus assay. First, a subsample of males and females were assayed immediately after feeding (”day zero”) to confirm virus viability. All fed males and females had detectable live infectious virus in their bodies, although females had statistically higher viral titers (*P* = 0.005), likely because they could physically ingest a larger volume of blood. At day 7 post-exposure, viral titers were not statistically different between males and females in the bodies, the legs/wings, or the saliva (Figure 6A). At day 14 post-exposure, females had higher viral titers in their bodies (*P* = 0.001) and legs/wings (*P* = 0.0083) compared to males, suggesting either greater viral replication rates, or simply more tissue available for virus replication due to larger size. However, viral titers in saliva between males and females did not differ statistically (Figure 6B).

**Figure 6.**
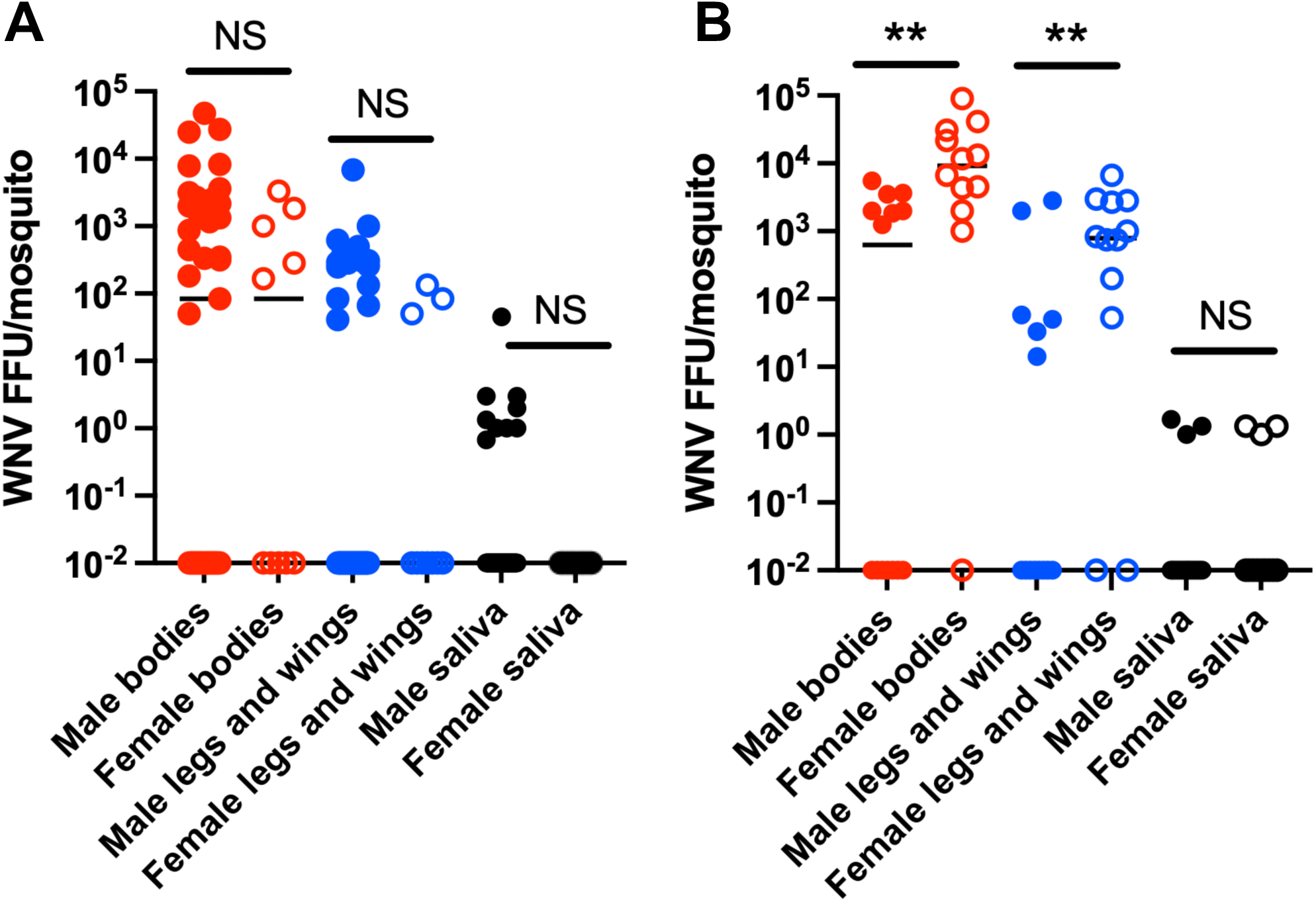
West Nile virus vector competence for male and female *Cx. tarsalis*. A) 7 days post-exposure. B) 14 days post-exposure. Red = bodies (infection); blue = legs/wings (dissemination); black = saliva (transmission). Males = closed circles, females = open circles. Zero values had 0.01 added purely for log-scale plotting purposes (10^-2^ = uninfected); analysis was performed on untransformed data. ** = *P* < 0.01.

### Vertebrate DNA and blood can be detected in wild-caught male mosquitoes

While BG-Sentinel traps are designed to collect female mosquitoes, they do also occasionally catch males [e.g. 20-21]. During collections in Texas, we collected a total of 15 male mosquitoes (ten *Ae. albopictus*, one *Ae. sollicitans,* and four *Cx. quinquefasciatus*). Vertebrate DNA was not detected in *Ae. albopictus* or *Ae. sollicitans*. However, three of the four collected *Cx. quinquefasciatus* males had detectable vertebrate DNA (Ct values ranging from 15.55 to 18.36) (Supplementary Table 1). When homogenized during extraction, the supernatants from these positive males were reddish-brown, possibly suggesting the presence of blood in the specimen. We sequenced the vertebrate COI sequences in these samples; all 3 hit canine (best hits - *Canis lupis familaris*, or domestic dog) (Supplementary file 1). In collections in Mallorca, Spain, we collected a male *Cx. pipiens* biotype *pipiens* mosquito with what appeared to be partially digested blood in its midgut (Figure 7). We sequenced the vertebrate mitochondrial 16S RNA gene sequences in this sample, which suggested the mosquito had fed on a human host (Supplementary file 1).

**Figure 7.**
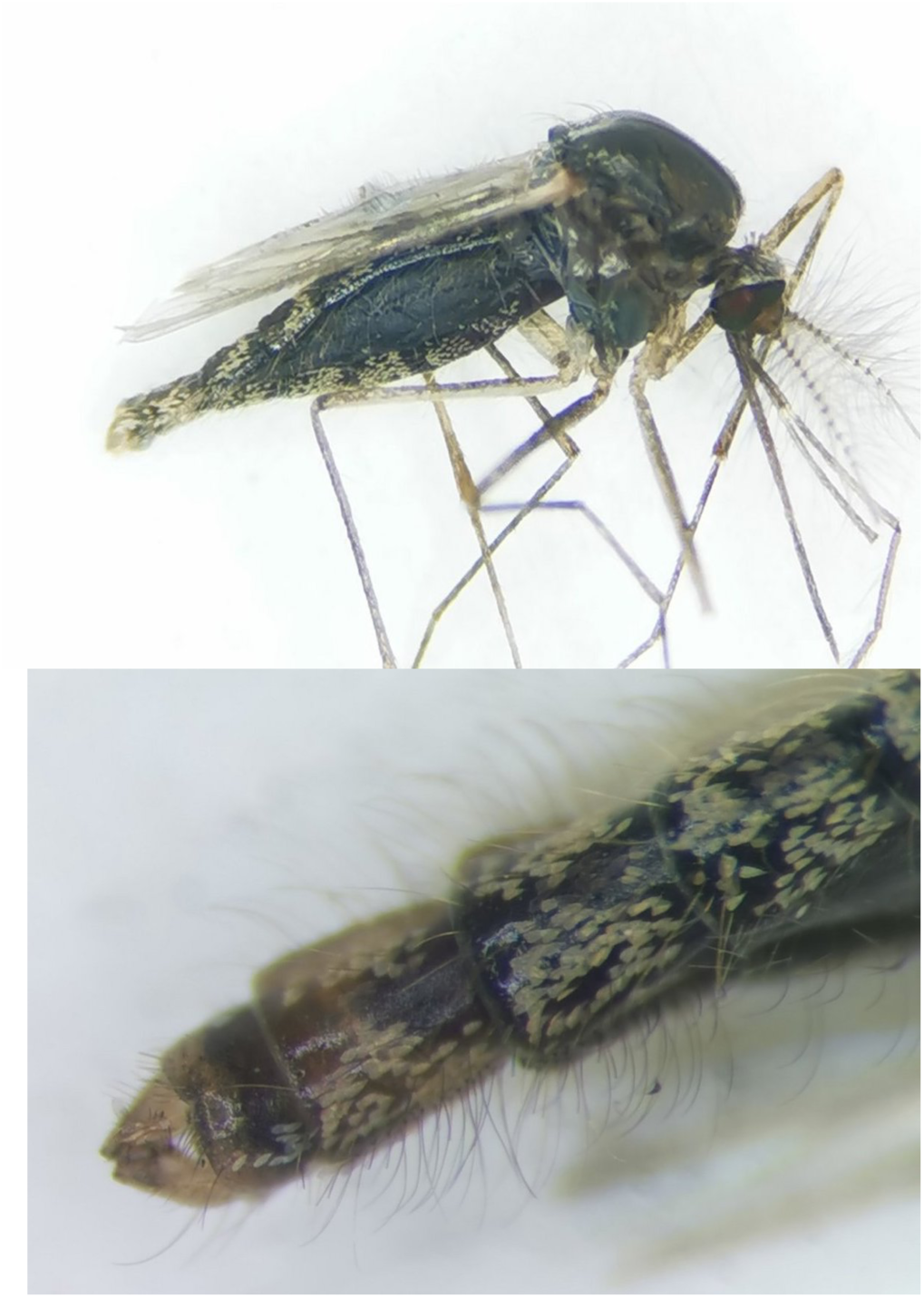
Wild bloodfed *Cx. quinquefasciatus* mosquito. Top: Male mosquito was collected incidentally during collections in Mallorca, Spain. What appears to be partially digested blood is visible in the midgut. Bottom: Closeup of mosquito genitalia to confirm sex of the specimen.

## Discussion

Previous work showed that in the laboratory, blood was toxic to male *Cx. quinquefasciatus* mosquitoes [7], suggesting that in this species male bloodfeeding seems to be a maladaptive trait. In our study, we demonstrate that males of multiple species can tolerate bloodfeeding, and that male bloodfeeding behavior is driven by water homeostasis during dehydration conditions. When mosquitoes cannot sense humidity due to Ir93a mutagenesis, dehydration does not increase blood seeking behavior. These results are consistent with the role of dehydration on bloodfeeding behavior in female mosquitoes, where dehydration can stimulate females to increase their bloodfeeding rates as well [17, 24–25] and thus may reflect an adaptive trait where mosquitoes (female or male) can maximize their water intake during drought or periods of low relative humidity if other sources (nectar or free water) are not available.

The mouthparts of male mosquitoes are thought to be physically incapable of penetrating vertebrate skin; however, our data suggest that this paradigm may be incomplete; in our experiments male mosquito mouthparts were proven adequate to pierce a Parafilm membrane, as well as vertebrate (human and mouse) skin. Dehydrated males were significantly attracted to and actively probed the hand of a human host, and one individual was even able to penetrate the outer epidermis, leading to a transitory immune reaction (Supplementary Video 10 and Figure 4G). As the saliva of males differs from that of females, lacking various proteins needed for immunomodulation and bloodmeal acquisition [26], and it is likely that very little saliva was transferred compared to the bite of a female mosquito, it is not surprising that the host immune reaction was mild and rapidly resolving. When allowed access to a wound, dehydrated *Cx. tarsalis* male mosquitoes readily probed the wound and one took a bloodmeal. As this experiment was facilitated by the fortuitous presence of a pre-existing wound on the hand of the senior author, it could not be deliberately repeated (as we were not allowed to make a deliberate wound due to IRB concerns). Further experiments using *An. stephensi* and a mouse model demonstrated that male mosquitoes can and will bite vertebrate hosts and can canulate capillaries to acquire blood. In aggregate, these experiments suggest that male mosquitoes do have the ability to take blood from vertebrate hosts under specific circumstances and that although not necessary, feeding may be facilitated by the presence of host wounds. According to fossil evidence, male mosquitoes are thought to once have had the ability to feed on vertebrate blood, and to have lost this ability over evolutionary time [27]. It is possible that the neural circuitry regulating host seeking and bloodfeeding behavior may still be conserved among male mosquitoes, or alternatively that this is simply a unique response to dehydration conditions to allow replenishment of water reserves, as a lack of humidity detection observed in Ir93a mutants muted the blood feeding phenotype. However, more in-depth functional neural studies are required to fully investigate this phenomenon.

Interestingly our data demonstrate that, if male *Cx. tarsalis* orally acquire a WNV infection, they are competent vectors and transmit the virus at similar rates and titers compared to females. In our experiments we explicitly used an assay that quantified live, infectious viral particles rather than quantitative PCR to rule out results that might be due to carryover of non-infectious viral RNA. Our results suggest that male *Cx. tarsalis* retain the receptors necessary for viral infection on their midgut, salivary glands, and other body tissues. Intriguingly, there is a recent report of wild-caught male *Cx. pipiens* mosquitoes that were infected with WNV from accidently feeding on infected blood droplets during a laboratory assay [28]. *Cx. tarsalis* males (as well as *Cx. pipiens* and other *Culex* mosquitoes) can also be infected with WNV due to vertical transmission from infected mothers [11, 29–30], and it is possible that the ability for males to become infected with virus orally is a consequence of their retaining receptors necessary for vertical infection.

Finally, there is the question “is bloodfeeding behavior by male mosquitoes epidemiologically significant?” It is already known that male mosquitoes can be indirectly important for vector-borne disease transmission dynamics. For example, mating can affect key physiological parameters in females related to pathogen infection and transmission [2]. More directly, in some species including *Cx. tarsalis* and WNV, male mosquitoes can be infected with arboviruses by vertical transmission from infected mothers [11, 29–30]. Infected males can also transmit some viruses venereally to females during mating where they can be transmitted to vertebrate hosts during feeding [29–30]. Consistent collection of males using host-derived attractants suggest that males are commonly found to move toward hosts [8], increasing the potential of male feeding on host-derived fluids under specific conditions (dry periods with a lack of water and/or sugar resources). In field collections, we were able to detect vertebrate DNA/blood in wild-caught *Culex* males in two separate geographic locations, suggesting that male mosquitoes do encounter vertebrate hosts for bloodfeeding, either through wounds or possibly by direct biting. In total, our study suggests the possibility of edge cases where male mosquitoes could be more directly implicated in pathogen transmission, where males undergoing dehydrating or starvation conditions (for example, during drought) acquire virus through vertical transmission from infected mothers or by feeding on an infected vertebrate host, then transmit to a naïve host through feeding on an open wound or by probing the skin, as mosquitoes often transmit the bulk of virus when probing skin prior to actually taking a bloodmeal [31].

We must emphasize that while compelling, the ability for male mosquitoes to bloodfeed or transmit pathogens in nature have not been widely studied (we suspect that researchers have not rigorously examined these phenomena as the dogma in the field is that males do not bloodfed). However, our data suggest that the canonical role of males as non-bloodfeeders needs to be re-examined and their contribution to pathogen transmission explicitly quantified (even if minor), particularly in light of recent vector-borne disease control strategies that rely on the mass release of male mosquitoes into natural populations [32–35].

## Supporting information

Supplementary File 1 - 16S and COI sequences

Sup video 1 - Cx tarsalis probing

Sup video 2 - Cx tarsalis probing

Sup video 3 - Ae aegypti probing

Sup video 4 - Ae aegypti probing

Sup video 5 - Ae notoscriptus probing

Sup video 6 - Ae notoscriptus probing

Sup video 7 - Cx quinquefasciatus probing

Sup video 8 - Cx quinquefasciatus probing

Sup video 9 - Ae aegypti landing behavior

Sup video 10 - Cx tarsalis biting behavior

Sup video 11 - Cx tarsalis wound feeding

## Declarations

### Ethics approval and consent to participate

All experiments with a human volunteer used JLR (PSU IRB Exempt Protocol STUDY00024284) or PAR (University of Melbourne Human Ethics committee approval 0723847). Mouse experiments were performed under Johns Hopkins University IACUC protocol MO21H10.

## Acknowledgements

This research was supported by NIH/NIAID grant R01AI150251, USDA Hatch Project 4769, a grant with the Pennsylvania Department of Health using Tobacco Settlement Funds, and funds from the Dorothy Foehr Huck and J. Lloyd Huck endowment to JLR, and NIH/NIAID grant R01AI148551 to JBB and JLR. RLMNA, MVB, DEN were supported in part by the Bloomberg Philanthropies; MVB was additionally supported by NIH/NIAID training grant 2T32AI138953-06A1. Mosquito collections in Spain were funded by the *Aedes* Invasive Mosquitoes AIM COST Action CA17108 funded by COST (European Cooperation in Science and Technology). PAR was supported by an Australian Research Council Discovery Early Career Researcher Award (DE230100067) funded by the Australian Government. RL was funded by the Federal Ministry of Research, Technology and Space of Germany (Grant Number 01Kl2022). We thank Ms. Francine McCullogh for laboratory assistance, Ms. Amelia Romo and Ms. Heather Engler for assistance with mosquito rearing, Mabel Tao and Cindy Kimura for assistance with preparing mice for mosquito feeds, Apeksha Warusawithana for assistance with probing experiments, and Paul Garrity for providing Ir93a mutant mosquitoes used in this study. Finally, we thank Jiji the cat for his participation in this research.

## Authors’ contributions

JB, REJ, RSK, AH, ML, PAR, RL, T-YC, MAM, CB, MAG, RG-L, RLMNA, MVB, DEN, JBB, and JLR conducted the research; JBB contributed materials and reagents; JB, REJ, PAR and JLR analyzed the data; JB, REJ, PAR, JBB, and JLR wrote the manuscript.

## Supplemental legends

**Supplementary Video 1.** Probing behavior of dehydrated male *Cx. tarsalis* mosquito on the thumb of a human volunteer.

**Supplementary Video 2.** Probing behavior of dehydrated male *Cx. tarsalis* mosquito on the index finger of a human volunteer.

**Supplementary Video 3.** Probing behavior of dehydrated male *Ae. aegypti* mosquito on the thumb of a human volunteer.

**Supplementary Video 4.** Probing behavior of dehydrated male *Ae. aegypti* mosquito on the index finger of a human volunteer.

**Supplementary Video 5.** Probing behavior of dehydrated male *Ae. notoscriptus* mosquito on the index finger of a human volunteer.

**Supplementary Video 6.** Probing behavior of dehydrated male *Ae. notoscriptus* mosquitoes on the index finger of a human volunteer.

**Supplementary Video 7.** Probing behavior of dehydrated male *Cx. quinquefasciatus* mosquito on the hand of a human volunteer.

**Supplementary Video 8.** Probing behavior of dehydrated male *Cx. quinquefasciatus* mosquito on the hand of a human volunteer.

**Supplementary Video 9.** Attraction and landing behavior of dehydrated male *Ae. aegypti* mosquitoes on the hand of a human volunteer.

**Supplementary Video 10.** Probing behavior of dehydrated *Cx. tarsalis* male mosquito on the wrist of a human volunteer. This mosquito succeeded in penetrating the outer epidermis.

**Supplementary Video 11.** Male *Cx. tarsalis* feeding from a wound on a human volunteer.

**Supplementary Table 1.** Ct values for vertebrate DNA detected in male mosquitoes collected from Galveston, TX.

**Supplementary File 1.** Vertebrate 16S and COI sequences obtained from field-collected mosquitoes, with associated BLAST statistics.

## Notes

### Competing Interest Statement

The authors have declared no competing interest.

### Summary of Updates

Added significant new laboratory and field data, along with multiple new co-authors.

