## Supplementary File 1 - 16S and COI sequences for "Revisiting the paradigm of anhematophagy in male mosquitoes"

**Supplementary File 1.** Vertebrate 16S and COI sequences obtained from field-collected mosquitoes.

**Mallorca Cxpipiens**

TTGAACCCATGTTTAACGGCCGCGGTACCCTAACCGTGCAAAGGTAGCATAATCACTTGTTCCTTAAATAGGGACCTGTATGAATGGCTCCACGAGGGTTCAGCTGTCTCTTACTTTTAACCAGTGAAATTGACCTGCCCGTGAAGAGGCGGGCATGACACAGCAAGACGAGAAACCCCGTATGGAGACTT

**Texas Cxquinqs1**

TACTTTATATTTACTGTTTGGAGCATGAGCCGGTACAGTAGGCACTGCTTTAAGCCTTCTAATTCGAGCCGAACTAGGTCAACCCGGTACCCTATTAGGAGATGACCAGATCTACAATCTAATCGTAACTGCCCATGCATTTGTAATAATTTTCTTTATAGTTATACCTATTATAATTGGGGCTTTGGAAACTGACTAGTTCCGCTAATAATTGGCACCCCAGACATGGCATTCCCTCGAATAAATAACATAAGTTTCTGATTGCTCCCCTCATCTTTTCTTCTCCTATTAGCGTCTTCTATGGTAGAAGCAGGTACAGGAACTGGATGAACTGTATACCCCTCACTAGCCGGCAACCTAGCCCACACAGGGGCATCAGTAGACCTAACAATTTTTTCCTTACATTTAGCTGGGGTCTCCTCCATTCTAGGGGCAATTAATTTTATCACTACTATTATTAATATAAAACCCCCAGCCATGTCCCAATACCAAACTCCCTTGTTTGTATGGTCCATACTAATTACAGCAGTCCTATTGCTATTATCATTGCCTGTATTAGCTGCTGGAATTACAATACTTCTGACAGACCAAAATCTAAACACAACATTTTTCGATCCTGCTGGAGGGGGAGACCCCATTCTATACCAACACTTATT

**Texas Cxquinqs3**

TACTCTATATTTACTATTCGGAGCATGAGCCGGTATAGTAGGCACTGCTCTAAGTCTTCTAATCCTAGCCGAACCGGGTCAACCTGATACCTTATTAGGGGATGACCAGATCTACAATGTAATCGTGACTGCCCATGCATTCGAATAATTTTCTTCATAGTCATGCCTATTATAATTGGGGGCTTTGGTAACTGACTAGTACCACTAATAATTGGCACCCCAGACATGGCATTCCCTTGAATAAATAATATAAGCTTCTGACTGCTTCCCCCATCCTTCCTTCTACTACTTGCATCTTCTATGGTGGAAGCAGGGGCAGGAACTGGATGAGCTGTATACACCCCACTAGCTGGCAACCTAACGGGGGCATCAGTAGACCTAACAATTTTCTCCTTGCATTTAGCAAGGGTCTCCTCTATTCTTGGTGCTATCAATTTTATCACCACTATTATTAATATAAAACCCCCAGCCATATCCCAATACCAAACTCCTTTATTTGTGTGATCCGTATTAATTACAGCAGTCCTACTATTATTATCATTGCCCGTACTAGCTGCCGGAATTACCATACTTTTGGCAGACCGAAATCTAAACACAACATTTTTCAACCCCGCTGGAGGGGGAGACCCCATCCTGTACCAGCACTTATT

**Texas Cxquinqs4**

CGAATGAACAACATGAGCTTCTGACTCCTTCCTCCATCCTTTCTTCTACTATTAGCATCTTCTATGGTAGAAGCAGGTGCAGGAACGGGATGAACCGTATACCCCCCACTGGCTGGCAATCTGGCCCATGCAGGAGCATCCGTTGACCTTACAATTTTCTCCTTACACTTAGCCGGAGTCTCTTCTATTTTAGGGGCAATTAATTTCATCACTACTATTATCAACATAAAACCCCCTGCAATATCCCAGTATCAAACTCCCCTGTTTGTATGATCAGTACTAATTACAGCAGTTCTACTCTTACTATCCCTGCCTGTACTGGCTGCTGGAATTACAATACTTTTAACAGACCGGAATCTTAATACAACATTTTTTGATCCCGCTGGAGGAGGAGACCCTATCCTATATCAACACCTATTCTGATTTTTCGGCCACCCAGAA

**Best hits:**

**Mallorca Cxpipiens**:

KJ446565.1

Homo sapiens isolate HGDP00552 mitochondrion, complete genome

97.33%

Ev = 6e-82

**Texas Cxquinqs1:**

MZ099455.1

Canis lupus isolate Wolf3-T-COI-1.seq cytochrome c oxidase subunit I (COI) gene, partial sequence; nuclear copy of mitochondrial gene

95.59%

Ev = 0.0

**Texas Cxquinqs3:**

MZ099455.1

Canis lupus isolate Wolf3-T-COI-1.seq cytochrome c oxidase subunit I (COI) gene, partial sequence; nuclear copy of mitochondrial gene

85.84%

Ev = 0.0

**Texas Cxquinqs4:**

PQ305541.1

Canis lupus familiaris voucher CN-HN-01 cytochrome c oxidase subunit I (COX1) gene, partial cds; mitochondrial

99.32%

Ev = 0.0
