## Supplementary Table 1 for "Revisiting the paradigm of anhematophagy in male mosquitoes"

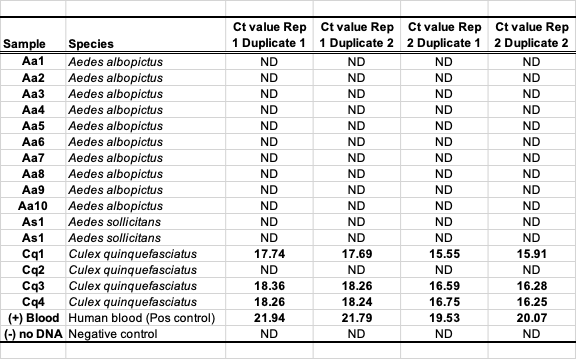
**Supplementary Table 1.** Ct values for vertebrate DNA detected in male mosquitoes collected from Galveston, TX.
